# Multicompartmental Non-invasive Sensing of Postprandial Lipemia in Humans with Multispectral Optoacoustic Tomography

**DOI:** 10.1101/2020.06.25.171413

**Authors:** Nikolina-Alexia Fasoula, Angelos Karlas, Michael Kallmayer, Anamaria Beatrice Milik, Jaroslav Pelisek, Hans-Henning Eckstein, Martin Klingenspor, Vasilis Ntziachristos

## Abstract

Disturbed blood lipid profiles after food intake have been associated with increased risk for cardiovascular and metabolic disease. Postprandial lipid profiling (PLP) can be used as a risk factor for cardiovascular disease, insulin resistance and other metabolic diseases but is based today on frequent blood sampling over several hours after a meal, an approach that is invasive and inconvenient for patients. Non-invasive PLP may offer a favorable alternative for disseminated monitoring in humans. In this study, we investigate the use of localized lipid sensing guided by Multispectral Optoacoustic Tomography (MSOT) for non-invasive, label-free assessment of postprandial lipemia in human vasculature and in soft tissues. For penetrating deep in human tissue, we utilize measurements at 930 nm, where lipids exhibit strong light absorption in the near-infrared spectral range (NIR). In a pilot study, we longitudinally measured postprandial lipemia in healthy subjects over 6 hours following consumption of a high-fat meal. Localized measurements were obtained from four anatomical structures: the radial artery, the cephalic vein, the brachioradialis muscle and the subcutaneous fat of the forearm. Analysis of optoacoustic signals demonstrated a 63.4% mean lipid increase in intra-arterial lipids at approximately 4 hours postprandially, a 89.7% mean increase in intra-venous lipids at 3-hours, a 120.8% mean increase in intra-muscular lipids at 3-hours and a 30.5% mean increase in subcutaneous fat lipids at 4-hours. We discuss how portable MSOT offers unprecedented potential to study lipid metabolism that could lead to novel diagnostics and prevention strategies by offering label-free and non-invasive detection of tissue biomarkers implicated in cardiometabolic diseases.

## Introduction

High blood lipid levels at either fasting or postprandial states indicate high risk for developing cardiovascular (CVD) and metabolic diseases, such as coronary artery disease (CAD), stroke, peripheral arterial disease (PAD), obesity, type 2 diabetes and non-alcoholic fatty liver disease (NAFLD) [1–7]. Although fasting blood lipid measurements have been successfully employed over the past decades for disease risk stratification [8, 9], the postprandial blood lipid levels are often better predictors of acute complications, such as heart attack, stroke or death [10]. For example, an analysis of the lipid profiles in 42710 patients at the non-fasting state showed that the non-fasting levels of lipids predicted increased risk of cardiovascular events [9]. Furthermore, lipid analysis of 8270 subjects provided strong evidence to support the routine use of postprandial lipid levels in clinical practice for accurate risk assessment of atherosclerotic CVD [11].

Assessment of fasting blood lipid concentrations is primarily done by single-time point blood sampling via venipuncture. Measurements of blood lipid dynamics can also be carried out by several blood samplings after the consumption of a high-fat meal to yield more information on lipid metabolism (postprandial lipid profiling, PLP) [12]. In both cases, the acquisition of blood samples causes patient discomfort, consumes hospital time and resources, and is only appropriate for infrequent sampling. Importantly, these methods of sampling lipids only allow observations in the blood and not in different tissue types. Recently, there has been interest in developing non-invasive lipid measurements to circumvent the need to obtain blood samples. Current non-invasive methods for lipid measurements in humans include breath measurements or eye image analysis. Lipid metabolism produces volatile organic compounds (VOCs, such as 2-pentyl nitrate, carbon dioxide, methyl nitrate, toluene etc.), some of which are exhaled. In one study, these VOCs were exploited to quantify blood lipid levels [13]. Nevertheless, breath analysis does not offer a direct measurement of lipids in blood or tissues and is vulnerable to time delays between changes in blood lipid levels and the corresponding changes in breath VOCs. Furthermore, the method was tested exclusively on subjects after an overnight fast without investigating possible influences on the measured VOC concentrations due to systemic metabolic changes triggered during the postprandial state. Lipids can also accrue in the cornea after arriving through the blood stream of the limbal vessels under hyperlipidemic conditions. Processing of images taken from the human eye (RGB colour representation) calculated the corneal lipid deposition by analyzing the intensity level within the region of lipid deposits [14]. Based on these values and known lipid values in the blood, a regression model was developed and used to estimate the blood lipid levels from the extracted image parameters. Similar to breath gas analysis, this approach only indirectly determines lipid concentrations in blood, based on the assumption of a relationship between lipid values in the corneal images and blood values. However, different physiological conditions that affect lipid accumulation in the cornea, including age, and experimental parameters that affect photographic fidelity may limit the estimation accuracy of this method. Both non-invasive methods described above provide estimates of lipid concentration in blood based on mathematical models that use an indirect correlation analysis with either measured VOCs or image intensities and have yet to be integrated into research protocols or clinical practice.

Novel techniques providing direct imaging of lipid distributions and dynamics in blood vessels and tissues could enable non-invasive tests that evaluate the risk for CVD and metabolic diseases, as well as the easy monitoring of nutritional and other metabolic conditions that are difficult to study. Here we aimed to introduce a method that could go beyond the current state of the art in measuring postprandial lipid dynamics by satisfying three critical specifications. First, the method should be safe, non-invasive and portable, so that it can be seamlessly disseminated to studies of large populations. Second, it should be capable of recording lipid measurements in different tissue compartments, not only single point bulk measurements, enabling differential studies of lipid circulation and uptake in tissues of interest. Third, it should be capable of frequent sampling to provide a detailed profile of highly resolved spatio-temporal lipid dynamics. Introducing such functionality into lipid research and medical care could significantly expand our knowledge of individual responses to nutritional challenges and offer new abilities for cardiovascular and metabolic risk assessment on a personalized basis.

To introduce this paradigm-shifting performance, we hypothesized that optoacoustic imaging, in particular multispectral optoacoustic tomography (MSOT), could offer a platform to obtain localized non-invasive sensing of lipid concentrations. MSOT acquires images of tissue molecules exhibiting optical absorption at multiple wavelengths in the near-infrared range (NIR) and resolves spectral information revealing deoxygenated hemoglobin (Hb), oxygenated hemoglobin (HbO_2_) and lipids among others (Fig. 1) [15]. Thus, MSOT should enable the monitoring of lipid distributions by means of wavelength selection within different tissue compartments. Preliminary evidence demonstrated that MSOT can resolve blood vessels, skeletal muscle and adipose tissues [16–21]. Furthermore, it was recently shown that MSOT can visualize oxidative metabolism by monitoring the rate of conversion of oxygenated haemoglobin to deoxygenated haemoglobin, revealing oxygen utilization by tissue [16, 22, 23]. More specifically, observations in animals were corroborated with measurements of oxidative metabolism during brown fat activation in humans and mice [16, 22, 23]. However, it is currently unknown whether MSOT can resolve lipid dynamics in response to nutritional inputs. Therefore, we employed MSOT to visualize vasculature and other soft tissue compartments in the 700-970 nm spectral window. Then, using observations at 930 nm, a wavelength where lipids exhibit an absorption peak in the near-infrared [24], we investigated whether we could resolve lipid-specific signals over time, in different tissue compartments, in response to oral fat intake. MSOT is a safe modality that uses light, does not require labels for imaging lipids, offers high resolution (< 300 μm) and large fields of view (> 4 cm) while reaching depths of 3-4 cm in living tissue. Each spectral image can be acquired in less than a second, which minimizes motion artefacts and allows for frequent longitudinal and comfortable measurements. We demonstrate in this study that these capabilities combine to afford a novel tool for the non-invasive study of lipid dynamics.

**Fig. 1.**
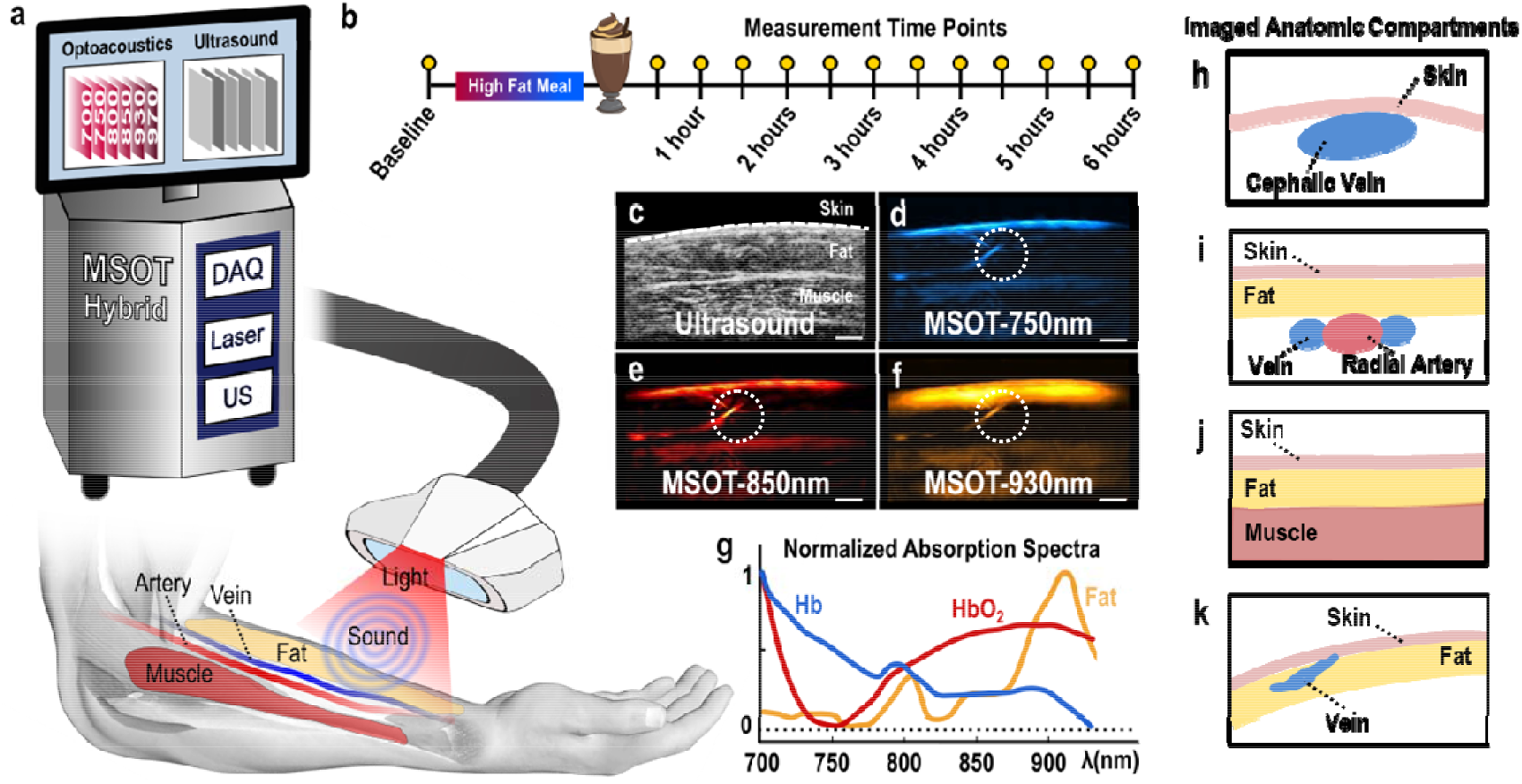
MSOT principle of operation and study design. (a) Configuration of the clinical hybrid MSOT/US system. (b) Postprandial lipemia measurement protocol. (c) Exemplary ultrasound image where the skin line (white dashed line), the subcutaneous fat and the skeletal muscle areas are shown. (d-f) MSOT images corresponding to the ultrasound image in (c). The dotted white circles show a small vessel detail within the subcutaneous fat region, demonstrating the excellent resolution performance of clinical MSOT technology. (d) MSOT image at 750 nm, representing mainly the distribution of deoxygenated Hb. (e) MSOT image at 850 nm, representing mainly the distribution of oxygenated Hb. (f) MSOT image at 930 nm, representing mainly the spatial distribution of lipids. Scale bars are 0.5 cm.(g) Absorption spectra in the near-infrared range (NIR) for Hb, HbO_2_ and fat/lipids. (h-k) Schematic diagrams of imaged anatomic compartments in the human forearm. (h) Cephalic vein. (i) Radial artery. (j) Skeletal muscle. (k) Subcutaneous fat.

## Methods

### Study design and experimental protocol

Four (n = 4) subjects (3 females and 1 male, age: 28 ±⍰7 years) with body mass indices (in kg/m^2^) of 28 (subject 1), 31 (subject 2), 23 (subject 3) and 21 (subject 4) were enrolled in the current pilot study. All volunteers were non-smokers and had no history of cardiovascular or metabolic disease. They were kindly requested to avoid consuming caffeine, food and alcohol for at least 12 hours before the planned measurements. All participants provided a written informed consent in full accordance with the work safety regulations of the Helmholtz Centre of Munich (Neuherberg, Germany).

Measurements took place in a dark, quiet room at a room temperature of ~23°C with the subjects in a sitting position. After a 12 h overnight fast, participants ingested an oral fat load within 5 min. The consistency of the fatty meal was 350 ml pasteurized heavy cream, 15 ml fat-free milk, 15 ml chocolate syrup and 1 tablespoon granulated sugar. This fat load contained 117 g fat (70 g saturated fat, 467 mg cholesterol), 41.5 g carbohydrate and 0.5 g protein and provided 1242 calories (86.4% from fat, 13.4% from carbohydrates, 0.2% from protein) [25]. The high-fat liquid meal was assigned to simulate the fat content of a typical high-fat meal (e.g. fast-food meal).

During the measurements, the probe was repeatedly placed in the same positions over the radial artery, the cephalic vein and the brachioradialis muscle of the dominant forearm, guided by stable skin markers (Fig. 1a). Recordings of these regions were taken post-fasting and then every 30 minutes and for 6 hours after consumption of the fatty meal (13 measurements in total), which was determined to be a suitable timespan to assess postprandial responses (Fig. 1b). The fourth participant felt some light gastric disturbances, so we decided to acquire less measurements and stop the experiment at 5 hours after the oral loading (5 measurements in total). Each anatomic compartment was scanned for ~10 sec.

Apart from prior anatomical knowledge, the arteries were initially identified from their pulsation, the veins from their compressibility with the hand-held probe, and the subcutaneous fat and the skeletal muscles by their characteristic textures in traditional ultrasound (US) (Fig. 1c). The identification of the different tissues and anatomical compartments was further facilitated by means of their MSOT appearance: the blood vessels and skeletal muscles were characterized by an increased absorption at the 750 nm and 850 nm, compared to the absorption at 930 nm, due to the strong presence of the Hb and HbO_2_ in these tissue compartments and the prominent absorption of both at these NIR-wavelengths [19]. Correspondingly, the subcutaneous fat tissue was characterized by an absorption peak at 930 nm where lipids absorb the most in the NIR (Fig. 1d-g).

### MSOT data acquisition

Measurements were conducted using a hybrid clinical MSOT/US (Acuity^©^, iThera Medical GmbH, Munich Germany). For ultrasound detection, the hand-held probe (Fig. 1a) was equipped with 256 piezoelectric elements with a central frequency of 4 MHz arranged in an arc of 145°. Illumination was achieved through an optical fiber, mounted on the same hand-held probe. Light was emitted in the form of short pulses (~10 ns in duration), at a rate of 25 Hz. For each pulse, almost 15 mJ of energy were delivered over a rectangle area of around 1 × 4 cm, ensuring compliance with the safety limits of laser use for medical applications [26]. For multispectral image acquisition, we employed 28 different light wavelengths (from 700 to 970 nm at steps of 10 nm). Thus, the recording of one ‘multispectral stack’, or a full set of 28 single-wavelength optoacoustic images, lasted ~1 sec. Co-registered US images were recorded in parallel to the MSOT images at frame rate of ~8 Hz.

As previously shown, MSOT images acquired at 750 nm reveal primarily Hb contrast, whereas images at 850 nm reveal contrast primarily from HbO_2_ [19]. MSOT images at the 930 nm show mainly tissue lipid distribution [16, 20, 24]. Thus, observed differences in the informational content, or else the appearance of different tissues, in the abovementioned single-wavelength MSOT images (Fig. 1d-f) are based on: i) the different features of the known absorption spectra of Hb, HbO_2_ and lipids in the NIR (Fig. 1g) [24] and ii) the content of Hb, HbO_2_ and lipids in the different tissues of each anatomic compartment.

The appearance of different tissues in the MSOT images (Fig.1d-f) showed good spatial correspondence to the co-registered US images (Fig.1c), but with additional functional and molecular contrast. Thus, illumination at 930 nm (Fig. 1f) highlights the subcutaneous fat region, which mainly contains lipids and is therefore characterized by much stronger light absorption or else optoacoustic image intensity, compared to adjacent blood vessels and muscles. The dotted white circles in Fig. 1d-f mark a small blood vessel in the subcutaneous fat region, highlighting the details that can be recorded by means of clinical MSOT, which achieves a spatial resolution of less than 300 μm.

Fig. 1h-k show the anatomical compartments, which were selected for analysis in MSOT images recorded at 930 nm in order to sense postprandial lipid dynamics in a variety of tissues: i) venous and arterial blood, the gold standard tissues employed for quantifying lipid dynamics in clinical practice, ii) skeletal muscle, which promotes the easy intra-tissue distribution of lipids via its high vascularization and the high-contrast blood lipid imaging due to its low lipid content under normal conditions (best-case scenario) and iii) subcutaneous fat, which is poorly vascularized compared to muscle, and provides a low-contrast environment for postprandial lipemia imaging due to its high lipid content (worst-case scenario).

### Data processing and analysis

Acquired MSOT data were reconstructed using a model-based reconstruction method [27]. The blood vessels (radial artery and cephalic vein) and the soft tissues of the forearm (subcutaneous fat and muscle) were first identified in consensus between two clinicians with experience in clinical MSOT and ultrasound imaging. The identification was based on anatomical knowledge, ultrasound guidance and the characteristic MSOT appearance for each tissue. For each subject, a set of characteristic 930 nm-frames for all measurements time points was selected. Next, precise regions of interest (ROIs) were manually segmented within the arterial or venous lumen, the subcutaneous fat and the muscle region. Finally, the mean intensity values of the pixels belonging to the manually-segmented ROIs were used to plot the time course of the MSOT-extracted lipid signals within each compartment during the lipemia challenge (*see Results*). Lipid signal calculations took place on the recorded 930 nm-images, without the application of any filtering or denoising.

## Results

Fig. 2 shows the MSOT imaging of postprandial lipid dynamics in the blood of veins (Fig.2a-d) and arteries (Fig.2e-h). Fig. 2a illustrates a series of characteristic MSOT images of a cross-section of the cephalic vein (subject 1) acquired at 930 nm that correspond to four time points: before oral loading (1), and 120 min (2), 240 min (3), and 330 min (4) after oral loading. The 3rd image (240 min after oral loading) corresponds to the time point of the maximum pixel intensities inside the venous lumen as measured by MSOT at 930 nm, and thus the maximum intravenous lipid content. Fig. 2b shows the pixel intensities along the profile lines of the previous image series (Fig. 2a). Fig. 2b provides insights into the contrast between the lipid signal detected inside the vascular lumen and the lipid signal of adjacent structures, which may be also affected during postprandial lipemia. The red regions in Fig. 2b show the range along the profile line (Fig. 2a) where the individual pixel intensity is above 50% of the maximum pixel intensity. The red regions serve as rough markers of the start and the end of the structure of interest (e.g. vascular lumen) along the profile lines. Our results show that both the spatial and temporal fluctuations in lipid signal inside the structures of interest are clearly higher compared to adjacent structures, proving that the observed phenomena are not caused by random fluctuations of the 930 nm-signal over the whole image, but by a postprandial increase in the blood lipid levels.

**Fig. 2.**
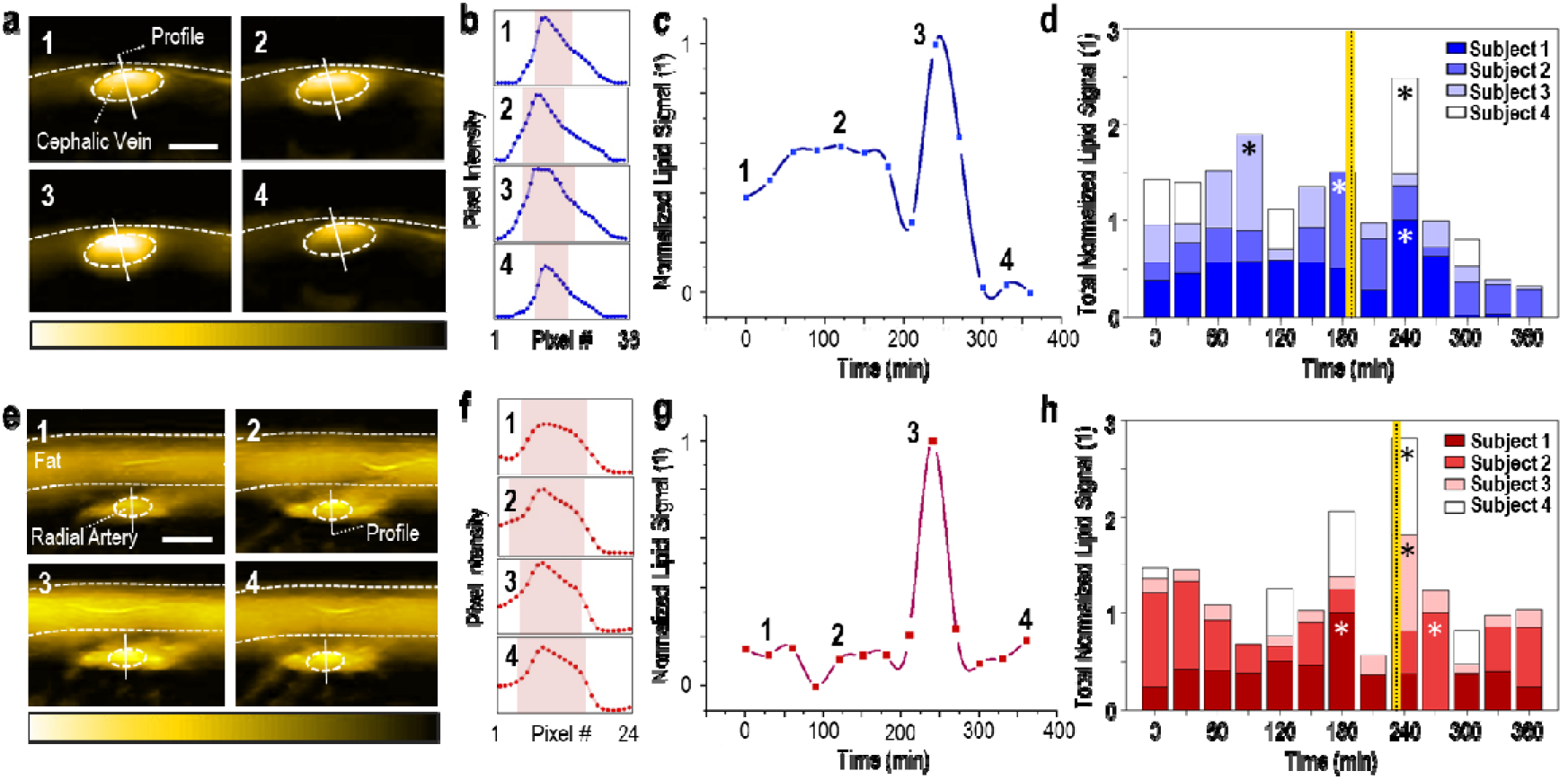
MSOT imaging of postprandial lipid dynamics in the blood of veins and arteries. (a) A series of cross-sectional MSOT images of the cephalic vein recorded at 930 nm, which correspond to the four time points indicated in (c) (subject 1). White dashed line: skin surface. White dashed ellipse: cephalic vein. Scale bars: 0.4 cm. (b) Pixel-intensity cross-sections along the corresponding profile lines in the image series of Fig. 2a. The red bands show the pixel range at 50% of the maximum pixel-intensity value along the profile line. (c) Normalized mean lipid signal within the cephalic vein for subject 1 during the whole postprandial lipemia test. The first time point corresponds to the fasting state. (d) MSOT-extracted lipid dynamics for the cephalic veins of all four subjects. The asterisks indicate the time points of the maximum recorded value for each subject. The vertical yellow-black line indicates the average time point (among all subjects) after oral loading for the maximum recorded lipid signal within the vein. (e) Cross-sectional MSOT images (at 930 nm) of the radial artery for the four time points of (g) (subject 3). Upper white dashed line: skin surface. Lower white dashed line: lower limit of the subcutaneous fat region. White dashed ellipse: radial artery. Scale bars: 0.3 cm. (f) Pixel-intensity cross-sections along the profile lines of Fig. 2e-image series. Red bands: pixel range at 50% of the maximum pixel-value along the profile line. (g) Postprandial lipid dynamics (normalized) within the radial artery for subject 3. (h) MSOT-extracted lipid dynamics for the radial arteries of all subjects. Asterisks: time point of the maximum recorded value for each subject. The vertical-black yellow line indicates the average recorded time point for the maximum lipid signal within the artery to be recorded.

Fig. 2c illustrates the changes in the mean lipid signals within the manually segmented ROI of the cephalic vein over the whole duration of the experiment for subject 1. The plotted mean values were normalized against the maximum mean value (at 240 min), with the numbered data points corresponding to the MSOT images in Fig. 2a. The first time point of Fig. 2c represents the baseline state before the oral fat loading. The observed lipid signal in the cephalic vein of subject 1 reaches its highest value (+ 27.6% compared to the subject’s baseline value) 240 minutes after the consumption of the high-fat meal (Fig. 2c, point 3).

Fig. 2d shows the normalized lipid signals recorded within the cephalic veins of all subjects for each predefined time point. For subject 2, the highest value was recorded at 180 minutes after oral loading (+ 111.4% compared to the baseline for subject 2). Readouts of subject 3 show a maximum lipid value of + 175.1% compared to same subjects’ baseline at 90 minutes after meal consumption. As already discussed, only five measurements were recorded for subject 4 over a 5-hour (300 min) observation time, due to light gastric disturbances (see Methods). The highest MSOT-measured lipid signal in the cephalic vein of subject 4 was recorded at 240 minutes postprandially (+ 44.5% compared to subject’s baseline). In summary, a mean maximum increase of + 89.7% was observed among the four subjects. The intra-venous lipid signal reaches its maximum value on average 187.5 minutes (~3 hours) after the oral fat loading. All percentages were extracted from the measured, and not the normalized, optoacoustic signal values.

Fig. 2e shows an exemplary set of lipids-MSOT images recorded at 930 nm of the radial artery cross-section for subject 3 at four different time points (30, 120, 240 and 360 minutes after fat loading). For subject 3, the intra-arterial lipid signal reaches its maximum (+69.3% compared to the subject’s baseline value) at 240 minutes (time point 3) after the consumption of the high-fat meal. Fig. 2f illustrates the intensity values for the pixels along the corresponding profile line for all images of Fig. 2e. Fig. 2g demonstrates the normalized mean lipid signal in the manually segmented arterial ROI for all predefined time points. As illustrated in Fig. 2h, which gives an overview of the radial artery lipid signals for all subjects, the highest intra-arterial lipid value for subject 1 was recorded at 180 minutes (+46.4%), for subject 2 at 270 minutes (+0.9%) and for subject 4 at 240 minutes (+136.9%) after the oral fat loading. To summarize, we observe a mean maximum increase of + 63.4%, with the highest intra-arterial lipid value being recorded at 232.5 minutes (~4 hours) after the oral loading on average. The herein presented results introduce MSOT as a new tool for *in vivo* tracking and quantification of lipid dynamics in human blood vessels at the postprandial state.

Fig. 3 shows the MSOT imaging of postprandial lipid profiles over time within soft tissues (skeletal muscle and the subcutaneous fat). Fig. 3a, presents a set of 930 nm-MSOT images of the brachioradialis muscle of the forearm taken from subject 2 at 60, 180, 270 and 360 minutes after the ingestion of the high-fat meal. The intra-muscular lipid signal reaches its highest level (+170.8% higher than the subject’s baseline) at 270 minutes postprandially. Fig. 3b depicts the pixel intensities along the profile lines of the image series presented in Fig. 3a. Fig. 3c shows the time course of mean lipid signals within the segmented muscle ROI during the whole test. Fig. 3d summarizes the MSOT-measured lipid dynamics within the skeletal muscles of all subjects: Maximum intra-muscular values were recorded at 210 minutes for subject 1 (+ 49%), 120 minutes for subject 3 (+ 145.5%) and 120 minutes for subject 4 (+ 117.7%). Thus, a mean maximum increase of + 120.8% is reported for the examined skeletal muscle. Finally, an average time span of 180 minutes (3 hours) after the consumption of the high-fat meal is needed for the lipid signal to reach its maximum within the forearm skeletal muscle ROI, as marked by the yellow-black vertical line in Fig. 3d.

**Fig. 3.**
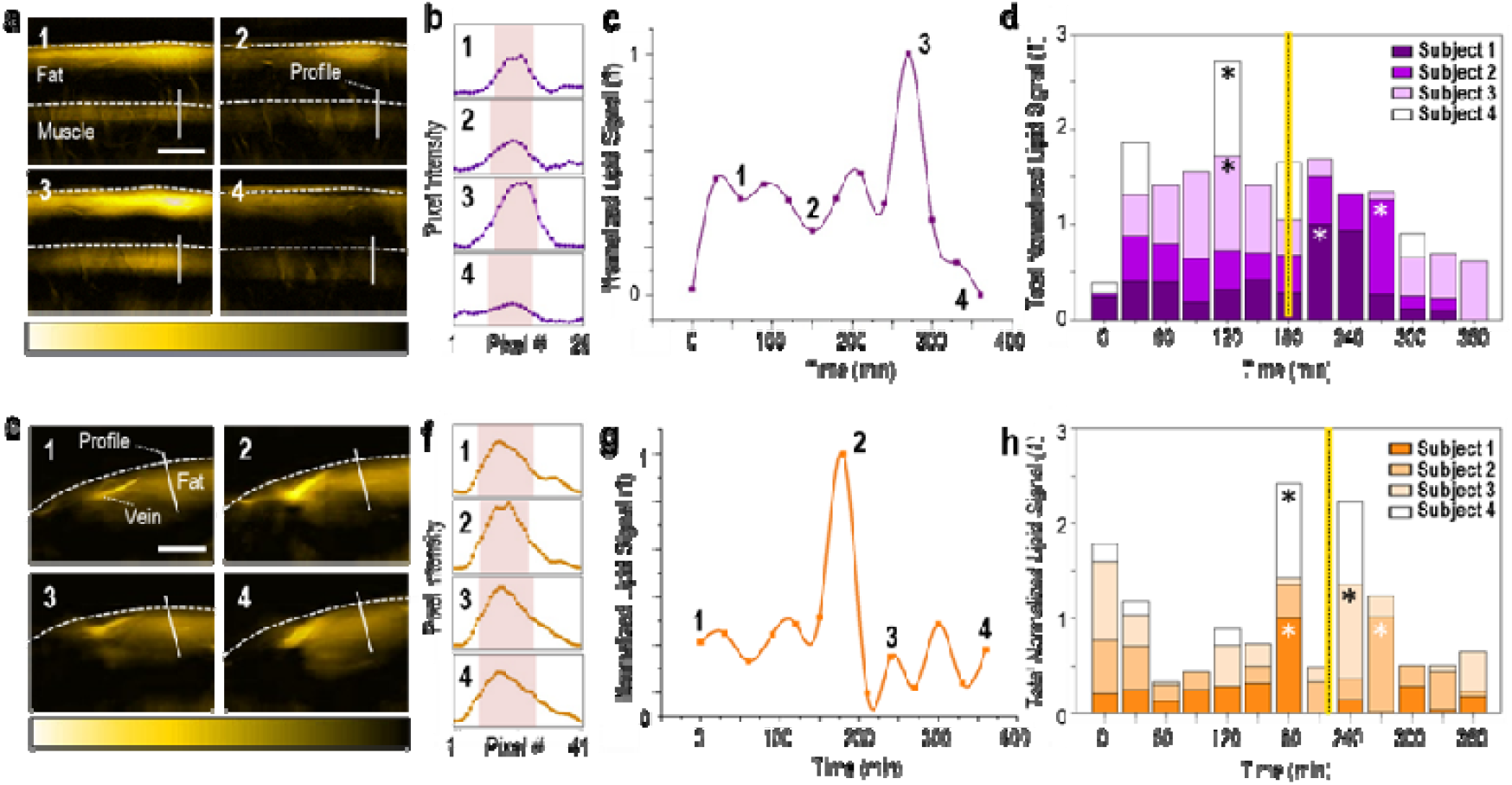
MSOT imaging of lipid dynamics in skeletal muscle and subcutaneous fat. (a) A series of cross-sectional MSOT images of the brachioradialis muscle recorded at 930 nm, which correspond to the four time points indicated in (c) (subject 2). Upper white dashed line: skin surface. Lower white dashed line: upper limit of the muscle region. Scale bars: 1 cm. (b) Pixel-intensity cross-sections along the corresponding profile lines in the image series of Fig. 3a. The red bands show the pixel range at 50% of the maximum pixel-intensity value along the profile line. (c) Normalized mean lipid signal within the muscle for subject 2 during the whole postprandial lipemia test. The first time point corresponds to the fasting state. (d) MSOT-extracted lipid dynamics for the brachioradialis muscles of all four subjects. The asterisks indicate the time points of the maximum recorded value for each subject. The vertical yellow-black line indicates the average time point (among all subjects) after oral loading for the maximum lipid signal within the muscle to be recorded. (e) Cross-sectional MSOT images (at 930 nm) of the forearm subcutaneous fat for the four time points of (g) (subject 1). Upper white dashed line: skin surface. Scale bars: 0.5 cm. (f) Pixel-intensity cross-sections along the profile lines of Fig. 3e-image series. Red bands: pixel range at 50% of the maximum pixel-value along the profile line. (g) Postprandial lipid dynamics (normalized) within the subcutaneous fat region for subject 1. (h) MSOT-extracted lipid dynamics for the subcutaneous fat of all subjects. Asterisks: time point of the maximum recorded value for each subject. The vertical-black yellow line indicates the average recorded time point for the maximum lipid signal within the subcutaneous fat to be recorded.

Fig.3e illustrates a time series of subcutaneous fat images from the forearm of subject 1 recorded at the 930 nm. The images were recorded at 0 (fasting state), 180, 240 and 360 min postprandially. Fig. 3f shows the pixel-intensities along the profile lines in the corresponding images in Fig. 3e. These images correspond to the time points 1-4 in Fig.3g, which presents the fluctuations of the normalized mean lipid signal inside the subcutaneous fat ROI for subject 1. Fig. 3h provides an overview of the lipid dynamics for all four subjects within the subcutaneous fat before and after the ingestion of the high-fat meal. Subject 1 shows a maximum increase of + 39.7% compared to the baseline at 180 minutes, subject 2 a maximum increase of + 25% at 270 minutes, subject 3 a maximum change of + 11.3% at 240 minutes and subject 4 a maximum increase of + 46.2% at 240 minutes postprandially. Thus, a mean maximum increase of + 30.5% in the lipid signal of the subcutaneous fat ROI is reported for the whole group. Moreover, this maximum lipid signal within the subcutaneous fat ROI was detected at 232.5 minutes (~4 hours) on average after the oral loading.

## Discussion

Postprandial lipemia is a dynamic condition characterized by a rise in blood lipid levels after the consumption of a meal, compared to the relatively stable lipid levels under fasting conditions. Pathological postprandial lipid profiles, in particular prolonged high lipid levels in blood, have been associated with serious diseases, such as diabetes, obesity and CVD [28]. Thus far, fluctuations of lipid levels in the blood stream have been monitored either by traditional blood sampling, or with non-invasive, but indirect, methods. We have demonstrated herein that clinical hand-held MSOT can non-invasively visualize and quantify lipid fluctuations in human blood vessels and soft tissues for several hours after the ingestion of a high-fat meal. This technique offers two key advantages over other non-invasive methods for *in vivo* lipid measurements: i) it provides direct lipid-specific molecular information in blood vessels and soft tissues without the need for injected contrast agents and ii) it has excellent spatial resolution of less than 300 μm for detailed tomographic imaging of *in vivo* lipid distributions over time. Our results agree with known postprandial lipid profiles from the literature [28] and show that MSOT is a viable alternative to blood sampling and appropriate for clinical applications.

The results of our pilot study demonstrate that MSOT can provide time-resolved data on lipid levels during the postprandial period. Light absorption by lipids reaches its peak in the NIR at 930 nm and signals recorded by MSOT in tissue upon illumination at this wavelength have been strongly associated with lipid content [24]. After consuming a fatty meal, our MSOT recordings at 930 nm showed a clear increase in signal intensities within the segmented anatomical compartments of interest (cephalic vein, radial artery, brachioradialis muscle and subcutaneous fat) approximately 3-4 hours on average after the subjects consumed a fatty meal. Our results agree with known patterns postprandial lipid fluctuations; significant increases in serum lipids have been recorded 3-4 hours after the ingestion of a high-fat meal using blood sampling [28]. Thus, our results demonstrate the unique capability of MSOT to provide direct and non-invasive monitoring of in *vivo* lipid level fluctuations at the postprandial state without the need for contrast agents. Moreover, although we only acquired data every 30 minutes for the current study, the high temporal resolution of MSOT (25 Hz) would enable the non-invasive investigation of faster phenomena of lipid kinetics to be explored in future studies.

MSOT also provided high resolution visualizations of lipids in blood vessels and soft tissues simultaneously, which is enabled by its 2-4 cm depth-penetration and approximately 4 cm horizontal field of view. Thus, using MSOT we were able to detect tissue variations in the occurrence of the highest average postprandial signals. The most intense signals in the cephalic vein and the skeletal muscle were recorded at ~3 hours while the corresponding signals in the radial artery and the subcutaneous fat were recorded at ~4 hours. To our knowledge, such rich and direct information on lipid dynamics has not been provided before in the literature. The ability to image both the blood circulation (arteries and veins) and the soft tissues (adipose tissue and muscle) during dynamic phenomena, such as the postprandial lipemia, may reveal metabolic interactions among the involved compartments and tissue components. Thus, MSOT could ideally help recognize paths that regulate the cross-talk between cardiovascular and metabolic components of human physiology and pathophysiology of diseases, such as obesity, hypertension and diabetes [29].

The current study introduces MSOT as a powerful tool for the non-invasive monitoring of *in vivo* blood lipid dynamics during the postprandial state. Our results open up new possibilities in the diagnostics and risk assessment of CVD and metabolic disease, especially when the high portability and low complexity of hand-held MSOT is expected to further facilitate its future disseminated use. Furthermore, the unique capability of MSOT technology to provide real-time and label-free visualizations of intra-vascular, intra-muscular and intra-subcutaneous fat lipid maps renders it an ideal tool for basic and clinical research in the cardio-metabolic field.

The imaging depth of MSOT (2-4 cm) is excellent compared to other optical techniques, but is limited compared to traditional clinical modalities, such as ultrasonography. Nevertheless, in our study we were able to access key blood vessels and soft tissues within these depth constraints and provide rich information on lipid dynamics in agreement with literature data. The consideration of novel light fluence correction models that compensate for intensity attenuation due to scattering and absorption is expected to further improve the precision of MSOT imaging deeper in tissue and widen the range of clinical applications. Furthermore, the development of phantoms and advanced spectral unmixing algorithms [30] will facilitate the direct quantification of lipid concentrations deep within muscle or other soft tissues, rather than only their relative fluctuations. Our aim here was to demonstrate a proof-of-concept via a human pilot study with a small number of healthy participants. More extended studies including larger cohorts of healthy volunteers and patients and simultaneous blood analyses are needed to further refine the application MSOT to the non-invasive monitoring of lipids.

Most individuals consume at least three meals per day and each meal is usually consumed before the postprandially high blood lipid levels return to baseline. Consequently, individuals are in a postprandial state for approximately 18 hours per day. Thus, the thorough investigation of *in vivo* lipid dynamics with novel methods may give new insights in several fields of basic and clinical cardio-metabolic research. It has been already shown that MSOT can provide precise anatomic, functional and molecular imaging of the vasculature and other soft tissues, such as adipose tissue and skeletal muscles [15, 16, 18, 19]. This unique set of capabilities may facilitate the exploration of hidden mechanisms of cardio-metabolic cross-talk by enabling multifaceted investigations of common cardiovascular and metabolic diseases, such as atherosclerosis, diabetes and lipid disorders. Further studies will advance hand-held MSOT toward its clinical translation with implications for objective diagnostics and therapy evaluation under patient-and operator-friendly conditions.

## Acknowledgements

We thank Dr. Robert Wilson for his attentive reading and improvements of the manuscript.

## Sources of funding

This project has received funding from the European Research Council (ERC) under the European Union’s Horizon 2020 research and innovation program under grant agreement No 694968 (PREMSOT) and was supported by the DZHK (German Centre for Cardiovascular Research) and by the BMBF (German Ministry of Education and Research) and by the Helmholtz Zentrum München, funding program “Physician Scientists for Groundbreaking Projects“.

## Disclosures

V. Ntziachristos has stock/stock options in iThera Medical GmbH. All other authors have no conflicts of interest to declare.

## Notes

### Competing Interest Statement

V. Ntziachristos has stock/stock options in iThera Medical GmbH. All other authors have no competing interests to declare.

